# Back Home: A Machine Learning Approach to Seashell Classification and Ecosystem Restoration

**DOI:** 10.1101/2025.01.08.632036

**Authors:** Alexander Valverde, Luis Solano

**Affiliations:** University of California, Santa Cruz; Joystick, FIFCO

## Abstract

In Costa Rica, an average of 5 tons of seashells are extracted from ecosystems annually. Confiscated seashells, cannot be returned to their ecosystems due to the lack of origin recognition. To address this issue, we developed a convolutional neural network (CNN) specifically for seashell identification. We built a dataset from scratch, consisting of approximately 19000 images from the Pacific and Caribbean coasts. Using this dataset, the model achieved a classification accuracy exceeding 85%.

The model has been integrated into a user-friendly application, which has classified over 36,000 seashells to date, delivering real-time results within 3 seconds per image. To further enhance the system’s accuracy, an anomaly detection mechanism was incorporated to filter out irrelevant or anomalous inputs, ensuring only valid seashell images are processed.

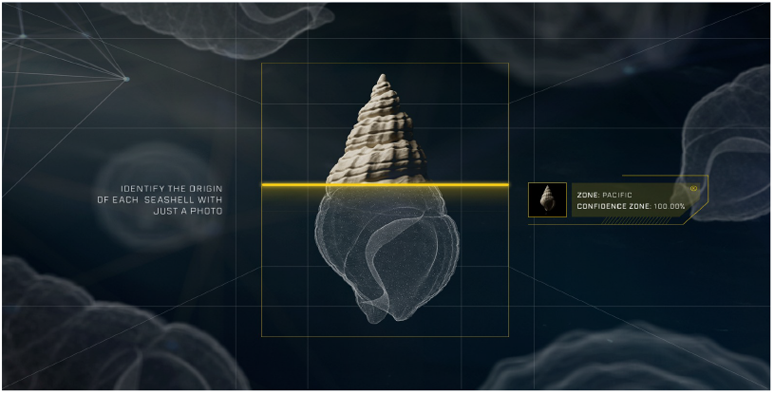

## 1 Introduction

Seashells are essential components of ecosystems, playing a vital role in maintaining ecological balance. As highlighted by Cheng et al. (2023) [3], seashells possess unique structural and functional characteristics, such as high hardness, toughness, corrosion resistance, and bioactivity, which make them crucial for various ecological and industrial applications. Many marine organisms rely on the work of seashells for their survival. Therefore, preserving seashells should be a matter of concern for everyone.

However, in recent years, a significant decline in seashells has been observed due to tourists collecting them from beaches in Costa Rica. Fortunately, these seashells are often confiscated at the Juan Santamaria Airport (Costa Rica’s national airport), but they cannot be returned because their exact ecosystem of origin (Pacific or Caribbean) is unknown.

To address this issue, this project developed a classification model capable of detecting the origin of each species from a single image. A total of 516 species registered in Costa Rica were analyzed, grouped into Pacific and Caribbean categories to perform binary classification. This resulted in a model suitable for returning confiscated seashells at the Juan Santamaria Airport to their respective ecosystems.

## 2 Related Work

Several studies have explored the classification of seashells and related marine objects, focusing on aspects such as feature extraction, classification methods, and dataset development. However, these approaches differ from our work, as they primarily emphasize individual feature extraction or taxonomic family classification rather than ecosystem-level insights.

Xue et al. (2021) addressed the challenge of identifying deep-sea debris using deep convolutional neural networks. They introduced the DDI dataset, comprising real deep-sea images, and developed the Shuffle-Xception network. This model incorporated innovative strategies, such as separable convolutions, group convolutions, and channel shuffling, to enhance classification accuracy [22]. Zhang et al. (2019) introduced a pioneering shell dataset containing 7,894 species with over 59,000 images, facilitating research in feature extraction and recognition. They applied traditional machine learning techniques, such as k-NN and random forest, to validate features like color, shape, and texture extracted from the dataset [23]. Yue et al. (2023) proposed “FLNet”, a convolutional neural network designed to address challenges in shellfish recognition, such as high feature similarity and imbalanced datasets. Their framework included innovative mechanisms for filter pruning and repairing to enhance feature representation, alongside a hybrid loss function tailored for unbalanced datasets. [24]

## 3 Our Work

The classification model was designed to address the critical challenge of determining the origin of confiscated seashells (Pacific or Caribbean). Due to the specificity of this task, we were required to build a comprehensive dataset from scratch, encompassing nearly every seashell species found in Costa Rica. This effort resulted in the creation of the first seashell database that integrates species from both the Pacific and Caribbean coasts of the country.

By leveraging advancements in convolutional neural network (CNN) architectures and integrating anomaly detection mechanisms, this model ensures high accuracy and reliability for real-world deployment. A user-friendly web application was also developed to facilitate the classification process (see Appendix A). This application enables users to upload images of confiscated seashells and obtain real-time predictions regarding their ecosystem of origin. The app was designed with simplicity in mind, ensuring accessibility for a wide range of users.

This section outlines the data preparation, model selection, and training strategies employed to achieve the desired goal.

### 3.1 Data Collection and Categorization

The dataset used for this research comprises 19,058 images representing 516 species of seashells, collected over 10 months. From the total of images, 9,553 are from the Caribbean and 9,505 from the Pacific. All of these species were categorized into four distinct groups based on their taxonomy and ecosystem (see Appendix B) To construct the dataset, we curated a comprehensive list of species by referencing specialized resources such as Conchology Inc. [11], iNaturalist [10], the Florida Institute of Technology [14], and ConchyliNet [4]. While other seashell datasets exist for image classification [13,23] they are not tailored to the unique biodiversity of our ecosystems. Therefore, we were required to source specific images for each species, capturing their distinct characteristics and possible variations to ensure accurate representation.

**Table 1:**
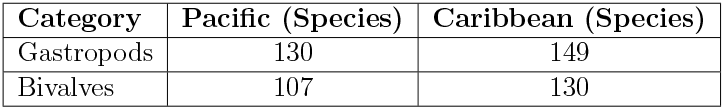
Comparison of species counts for Gastropods and Bivalves in the Pacific and Caribbean regions.

This categorization resulted in a total of 279 species from the Caribbean and 237 species from the Pacific. Each seashell was grouped by family, genus, and species to ensure accurate taxonomic representation. A detailed list of all categorized species is provided in the Appendix for reference. The dataset was meticulously verified to ensure the quality and consistency of the images, incorporating diverse positions, backgrounds, and environmental conditions to enhance the model’s robustness.

**Figure 1:**
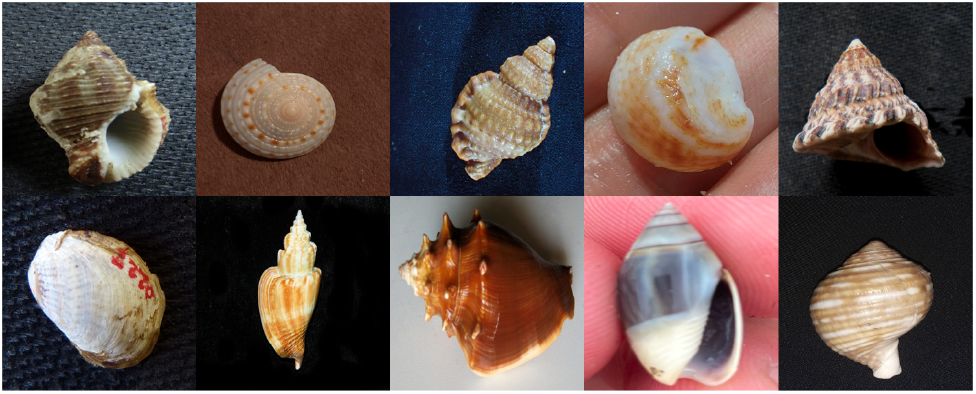
Representative Pacific seashells from the dataset.

**Figure 2:**
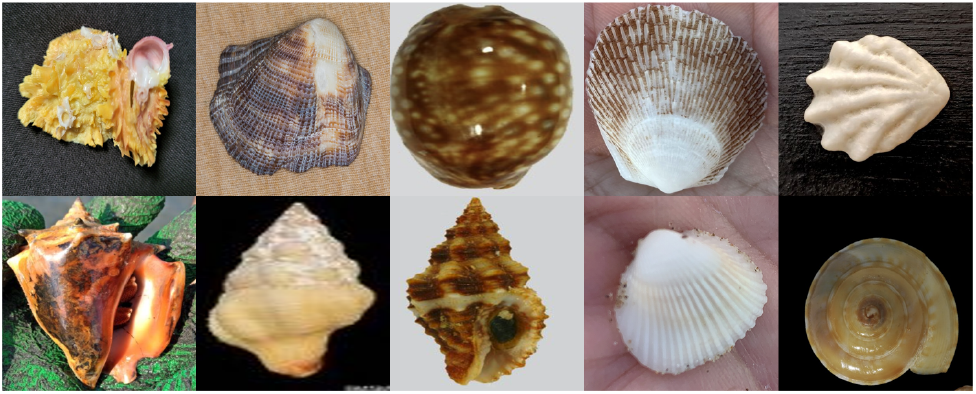
Representative Caribbean seashells from the dataset.

The dataset was divided into three subsets for training, validation, and testing, with 70% allocated to the training set for model training, 15% to the validation set for hyperparameter tuning and performance evaluation during training, and 15% to the test set for evaluating the final model performance on unseen data. This split ensured a balanced representation of families across all subsets, supporting a rigorous and reliable evaluation of the model’s performance. All images were resized to 224×224 pixels to maintain consistent input dimensions for the model architecture, while preserving key visual features necessary for classification.

To enhance the model’s ability to generalize and handle variations in real-world data, several data augmentation techniques were implemented. These included random rotations (±20 degrees), horizontal flips, and subtle adjustments to brightness and contrast. Additionally, random cropping and zoom operations were applied to simulate different viewing distances and perspectives. These augmentation techniques effectively expanded the training dataset and helped the model become more robust to natural variations in species appearances across different lighting conditions and viewing angles. These categorizations were critical in training the final classification model, as they allowed the system to learn distinctions not only between seashells but also between ecosystems. By leveraging this structured grouping, the model could focus on subtle morphological, color, and texture variations that characterize species from different ecosystems.

### 3.2 Model Training

The final classification model was built using the ConvNext architecture [12]. We deliberately excluded Vision Transformers (ViT) from our experimentation due to their substantially larger model size and computational requirements, which would have hindered deployment in resource-constrained environments. We experimented with several established architectures including ResNet50 [7], DenseNet121 [8], and MobileNetV2 [18]. Despite extensive training, these models consistently achieved accuracy scores below 81%, even when using identical training configurations (Table 5 - Appendix A).

ConvNext demonstrated superior performance, which we attribute to its 7 *×* 7 kernel design that effectively captures global features without relying on computationally expensive attention mechanisms. This choice not only improved accuracy but also reduced the model’s computational overhead, making it more scalable when deployed for many simultaneous users as we did during the classification process.

To train the model we used the ConvNext-Tiny version from PyTorch library with ImageNet-1K weights. The following hyperparameters were used: The learning rate was set to 0.01, and the optimizer used was Stochastic Gradient Descent (SGD) with a momentum of 0.9 and a weight decay of 0.001. The batch size was 16, and the model was trained for 100 epochs. Starting from epoch 65, the learning rate was decayed by a factor of *e*^−0.05^. Thirty layers were unfrozen during training, allowing for fine-tuning of the deeper layers, while a final dropout layer with a rate of 0.3 was applied to reduce overfitting.

**Figure 3:**
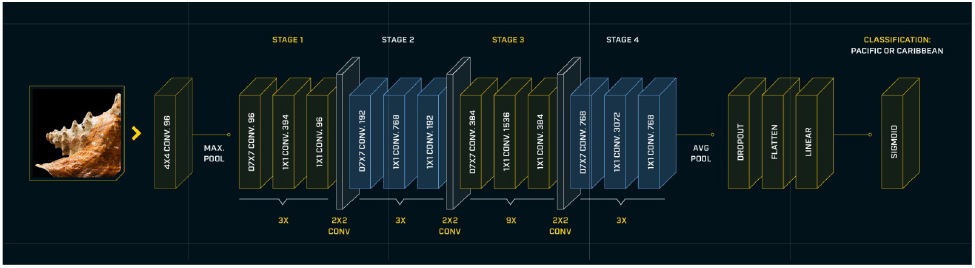
Classification process utilizing a modified ConvNext Tiny architecture

### 3.3 Anomaly Detection System

Anomaly systems have been broadly studied over recent years, especially for Large Language Models (LLMs). They have shown promising results in filtering out inputs that differ substantially from labeled datasets used for training [19–21].

Image analysis in real-world scenarios often requires systems that can identify when an input cannot be reliably assigned to any of the known classes, which is crucial in classification or detection tasks deployed in practical settings. Deep learning has emerged as a key enabler for such anomaly detection systems, such as Autoencoders and Generative Adversarial Networks to learn latent representations of normal data. By modeling what is typical, these methods can highlight inputs that fall outside the learned distribution. [1, 17]

Other approaches employ neural networks to estimate elements such as depth, perform feature extraction, or localize specific regions within an image, thereby providing a representation of the anomalous portion [2, 15]. Also, certain approaches emphasize feature extraction and are thus more aligned with our goal: to obtain a robust image representation and subsequently determine whether an input is anomalous relative to a trained reference dataset. [5, 6, 16]

Our approach focuses on using a vector representation of each training image and storing these vectors in a vector database. This enables efficient comparison against images provided by volunteers, resulting in a more accurate classification based on their similarities. This method ensures that only images containing the seashell as the primary element of interest are selected, reducing the impact of extraneous noise that could adversely affect the classification. Such an approach is particularly beneficial for a web application utilized by a wide range of users, many of whom may lack professional or scientific photography skills.

This system identifies images that do not correspond to seashells, preventing incorrect classifications during the real-time volunteering. The process involves creating image embeddings using a convolutional neural network (SqueezeNet1.0) [9]. Each image was transformed into a vector of 1,000 scalar values, representing a compact and discriminative feature set.

The embeddings were stored in a vector database to facilitate efficient similarity comparisons. When a new image is provided, its embedding is generated and compared against the stored embeddings using cosine similarity. The anomaly detection system operates as follows:

1. Compute the cosine similarity between the new image embedding and the top *k* closest embeddings in the database.
2. Average the similarity scores. If the resulting score is below a threshold (*λ*), the image is classified as an anomaly.

**Table 2:** Anomaly Detection Performance Across Object Categories (n=10 images per category except seashells with n=40). * Threshold score = 0.955. Images with mean similarity scores below threshold are classified as anomalies.

| Category | Images Below Threshold* |
| --- | --- |
| Cats | 10/10 |
| People | 7/10 |
| Buildings | 8/10 |
| Cars | 10/10 |
| Trees | 10/10 |
| Rooms | 9/10 |
| Cows | 9/10 |
| Hospitals | 8/10 |
| Horses | 9/10 |
| Dogs | 10/10 |
| Backgrounds | 10/10 |
| Ships | 9/10 |
| Birds | 9/10 |
| Frogs | 7/10 |
| Trucks | 10/10 |
| Airplanes | 9/10 |
| Reptiles | 6/10 |
| Electronic Devices | 9/10 |
| Insects | 8/10 |
| Seashells | 0/40 |

The cosine similarity between two embeddings, *u* and *v*, is defined as:

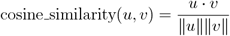

For anomaly detection, let *E*_input_ represent the embedding of the input image and {*E*_1_, *E*_2_, …, *E*_*k*_} represent the embeddings of the top *k* closest images in the database. The average similarity score (*S*) is calculated as:

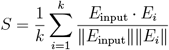

If *S < λ*, where *λ* is a predefined threshold, the image is classified as an anomaly:

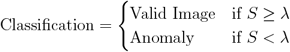

This method ensures that only valid seashell images are processed by the classification system, improving overall accuracy and reliability. The threshold *λ* was determined empirically by analyzing the distribution of similarity scores between known seashell images in our dataset. By computing clusters based on the similarities between images within the same class (either Pacific or Caribbean), we found that legitimate seashell images typically exhibited similarity scores above 0.955, which we established as our anomaly threshold.

**Figure 4:**
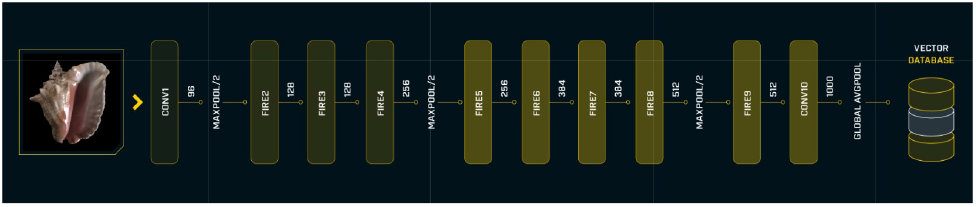
Embedding generation from images using the SqueezeNet architecture

## 4 Results

The classification model was evaluated on a test dataset containing 2,300 images, demonstrating its effectiveness in identifying the origin (Pacific or Caribbean) of seashells. The results show a general accuracy of 86.28%, with slight variations between the two locations. These metrics highlight the robustness of the model in real-world applications.

### 4.1 Results by Location

The performance of the model for each location is as follows:

- **Pacific**: The model achieved an accuracy of 85.43%, demonstrating strong performance in classifying seashells from the Pacific coast.
- **Caribbean**: The model achieved an accuracy of 87.10%, showing slightly better performance compared to the Pacific classification.

**Table 3:** Performance Metrics on Test Dataset (3,363 Images)

| Location | Accuracy (%) | Precision (%) | Recall (%) | F1 (%) |
| --- | --- | --- | --- | --- |
| Pacific | 85.43 | 88.70 | 85.43 | 87.00 |
| Caribbean | 87.10 | 83.45 | 87.10 | 85.29 |
| **Overall** | 86.28 | 86.23 | 86.13 | 86.17 |

The results highlight the model’s reliability in classifying seashells from both the Pacific and Caribbean coasts of Costa Rica. The slight performance variation between the two locations may be attributed to differences in image diversity, environmental factors, or species representation in the dataset. Even, with that accuracies, the model can be used in productive environments with security, as can be able to classify correctly 8 of 10 seashells,

## 5 Future Work

Future research should be focus on extending the model to other ecosystems by using the already trained weights. This approach could facilitate adaptation to new environments, improving the model’s versatility while reducing the need for extensive retraining. Additionally, enhancing the anomaly detection system for robustness and adaptability to diverse input conditions remains a promising direction.

## 6 Conclusions

This research successfully developed a robust classification model capable of identifying the origin of seashells from both Costa Rica’s Pacific and Caribbean coasts, effectively handling specimens from 515 different families. The results highlight the potential of machine learning to support ecological restoration efforts by facilitating the proper return of confiscated seashells to their native ecosystems.

In addition, we present a new dataset focus on seashell recognition from Pacific and Caribbean that can be used in future scientific research, we set a benchmark of 86.28% on the test dataset, that hopefully can be surpass by other scientists. This dataset not only served as the foundation for model development, but also represents a valuable resource for various institutions, enabling further research and advancements in marine biology and conservation.

Furthermore, the incorporation of an anomaly detection mechanism significantly enhanced the classification system’s reliability. Through effective filtering of anomalous and invalid inputs, the system maintained high accuracy levels and exhibited improved robustness when deployed in real-world environments. This implementation underscores the importance of developing systems capable of identifying and filtering out low-quality data before it affects the performance of production models.

## 7 Acknowledgements

We gratefully acknowledge the funding and help provided by FIFCO and its brand, Imperial. Their contribution was very important during this research and ensured its successful completion. Additionally, we wish to thank Dr. Yolanda Camacho for her invaluable work and dedication in compiling the lists of seashells. We also express our gratitude to the Sistema Nacional de Áreas de Conservación (SINAC), Universidad de Costa Rica (UCR), and AERIS for their support throughout this process.

## A Web Application Details

To make the seashell classification model accessible to a broad audience, a user-friendly web application was developed. This application allows users to upload images of confiscated seashells and receive real-time predictions regarding their origin (Pacific or Caribbean).

The interface was designed to ensure usability for non-experts, enabling seamless interaction and accurate results within seconds. The application integrates the classification model and anomaly detection system described earlier in this document, providing a robust platform for real-world use. It supports images captured in various conditions, ensuring flexibility and reliability. The key features of the application include:

- Image Upload: Users can upload single or multiple images of confiscated seashells.
- Real-Time Classification: The app delivers predictions in under three seconds per image, indicating the ecosystem of origin (Pacific or Caribbean).
- User-Friendly Design: The interface is intuitive and accessible to a wide range of users, requiring no prior technical expertise.

**Figure 5:**
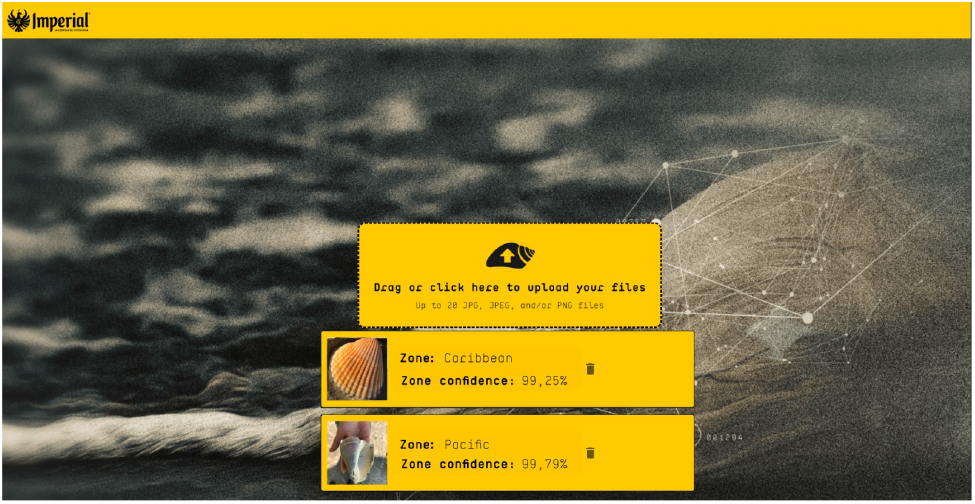
The web application for seashell classification, enabling real-time predictions.

**Table 4:** Summary of Model Results and Parameters.

| Model Version | unfreeze_layers | Optimizer | learning_rate | num_epochs | lr_epoch | Test Accuracy (%) |
| --- | --- | --- | --- | --- | --- | --- |
| ConvNext Tiny | 0 | SGD | 0.01 | 50 | 25 | 83.24 |
| ConvNext Tiny | 0 | SGD | 0.01 | 50 | 25 | 81.11 |
| ConvNext Tiny | 10 | SGD | 0.01 | 100 | 50 | 80.15 |
| ConvNext Tiny | 10 | SGD | 0.01 | 150 | 75 | 84.56 |
| ConvNext Tiny | 10 | SGD | 0.01 | 250 | 125 | 84.16 |
| ConvNext Tiny | 13 | SGD | 0.01 | 150 | 75 | 84.16 |
| ConvNext Tiny | 30 | SGD | 0.01 | 100 | 65 | 86.28 |

The **Test Accuracy (%)** column reflects the model’s performance on the test dataset. Notably, the final version of the model achieved the highest accuracy of 86.28%, obtained by unfreezing 30 layers, training for 100 epochs, and applying a learning rate decay starting at epoch 65.

**Table 5:** Accuracy scores (mean ± standard deviation) for models over 50 runs.

| Architecture | Accuracy (%) |
| --- | --- |
| ResNet50 | $78.3 \pm 0.4$ |
| DenseNet121 | $80.2 \pm 0.3$ |
| MobileNetV2 | $79.3 \pm 0.5$ |

## B List of Species

**Table 6:**
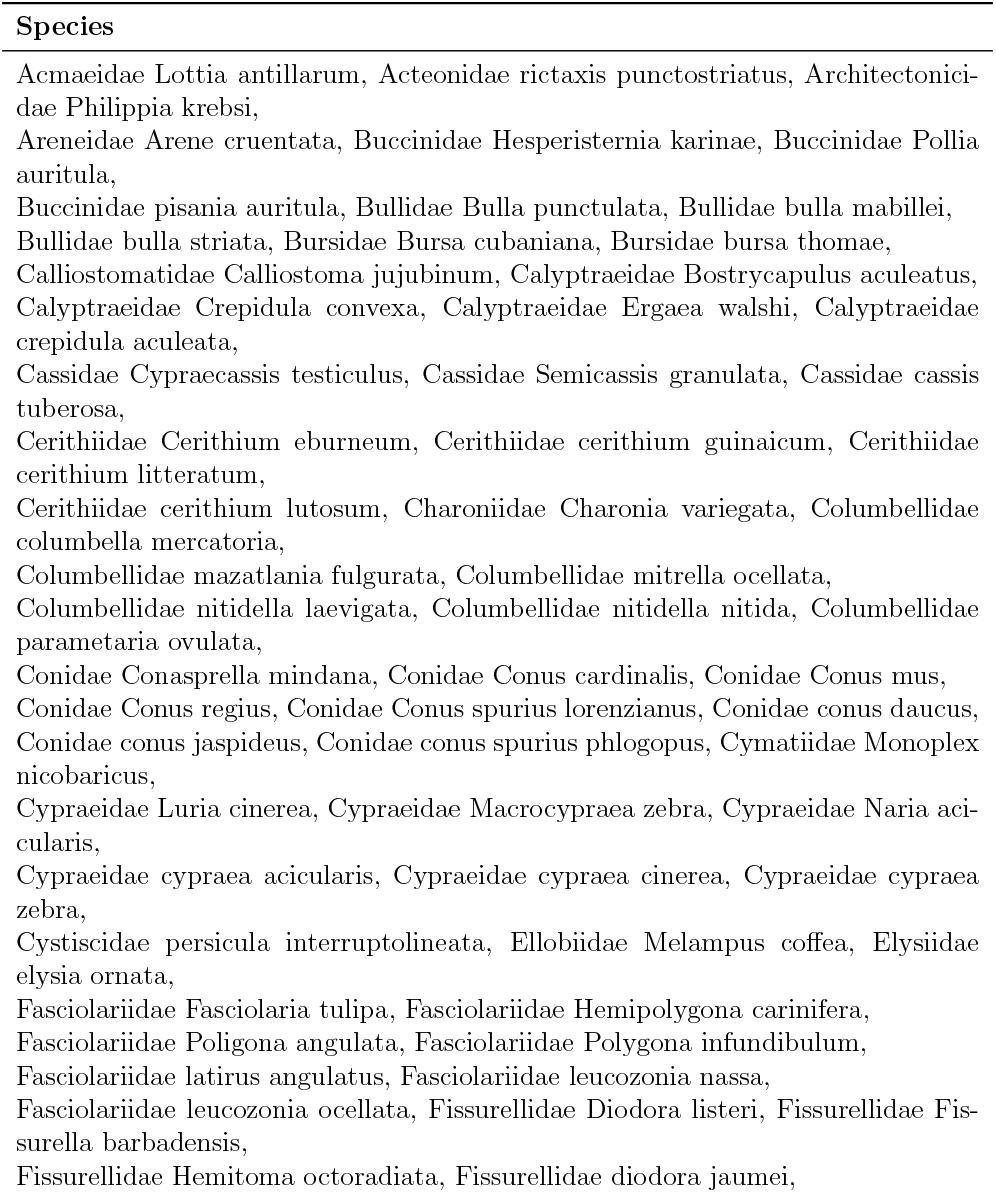
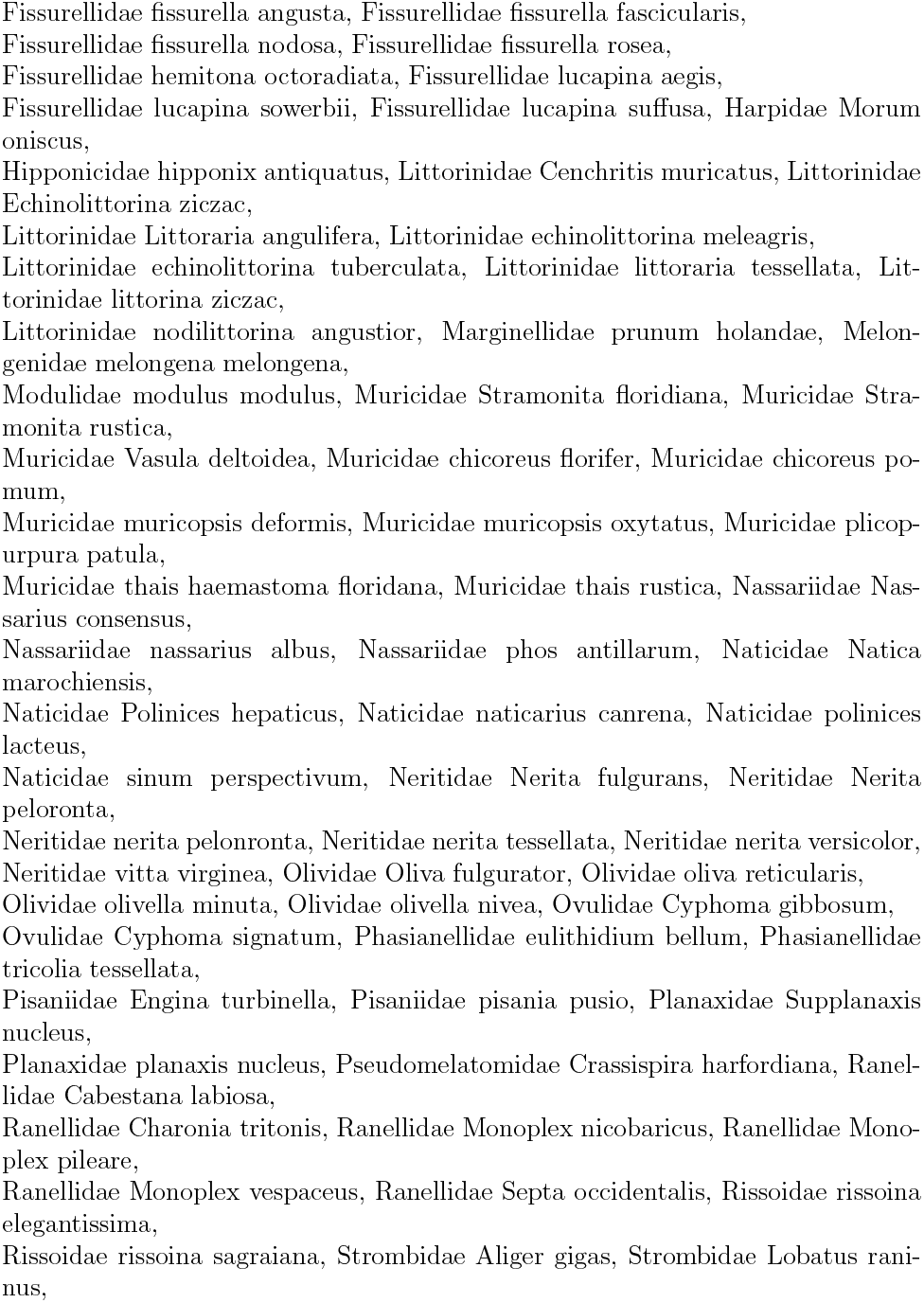
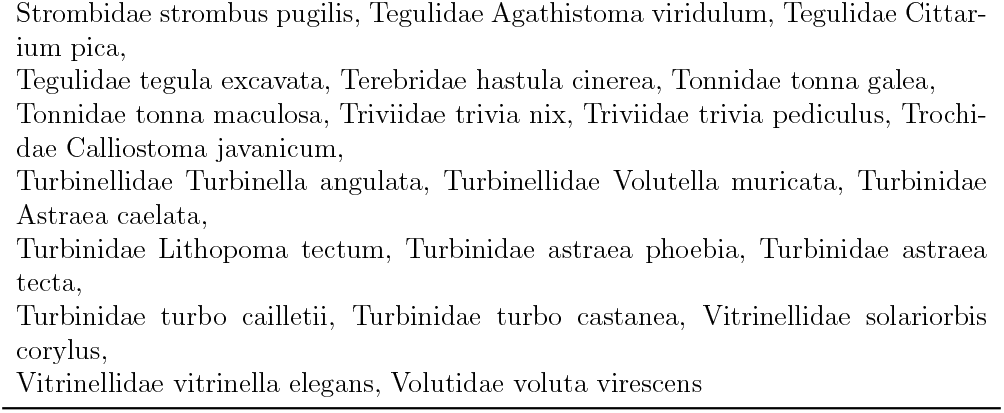
Seashell Species - Caribbean Gastropoda.

**Table 7:**
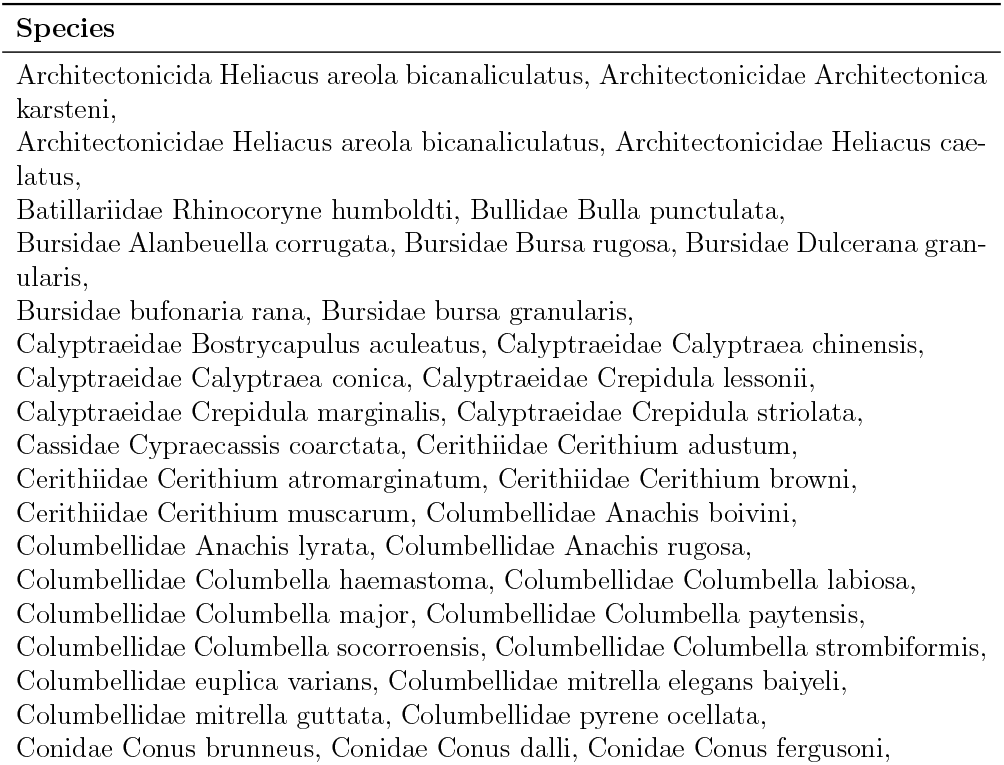
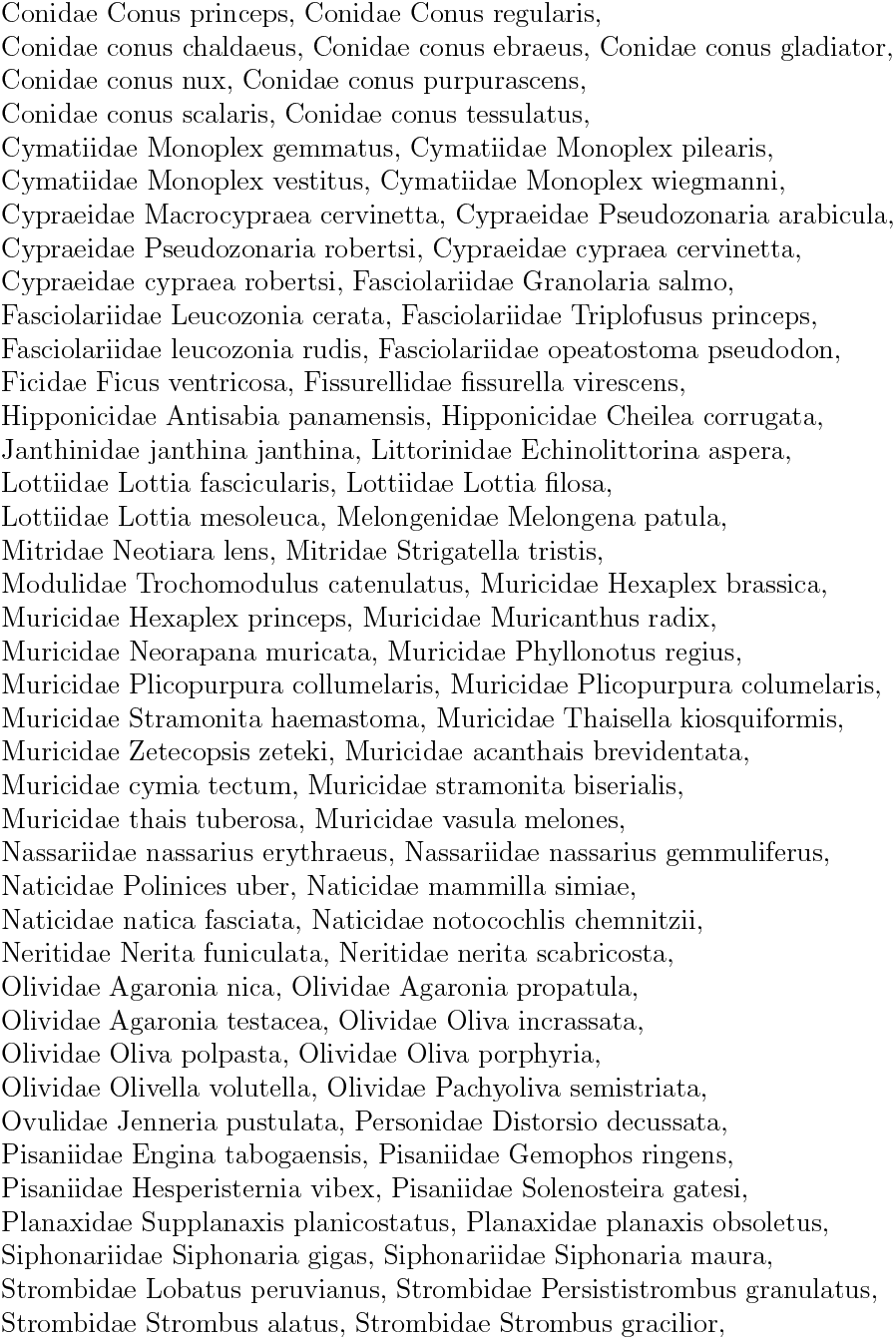
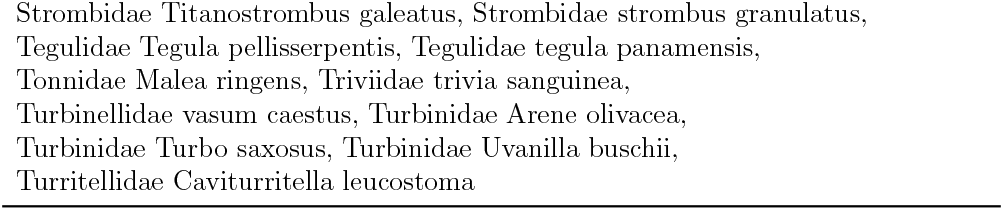
Seashell Species - Pacific Gastropoda.

**Table 8:**
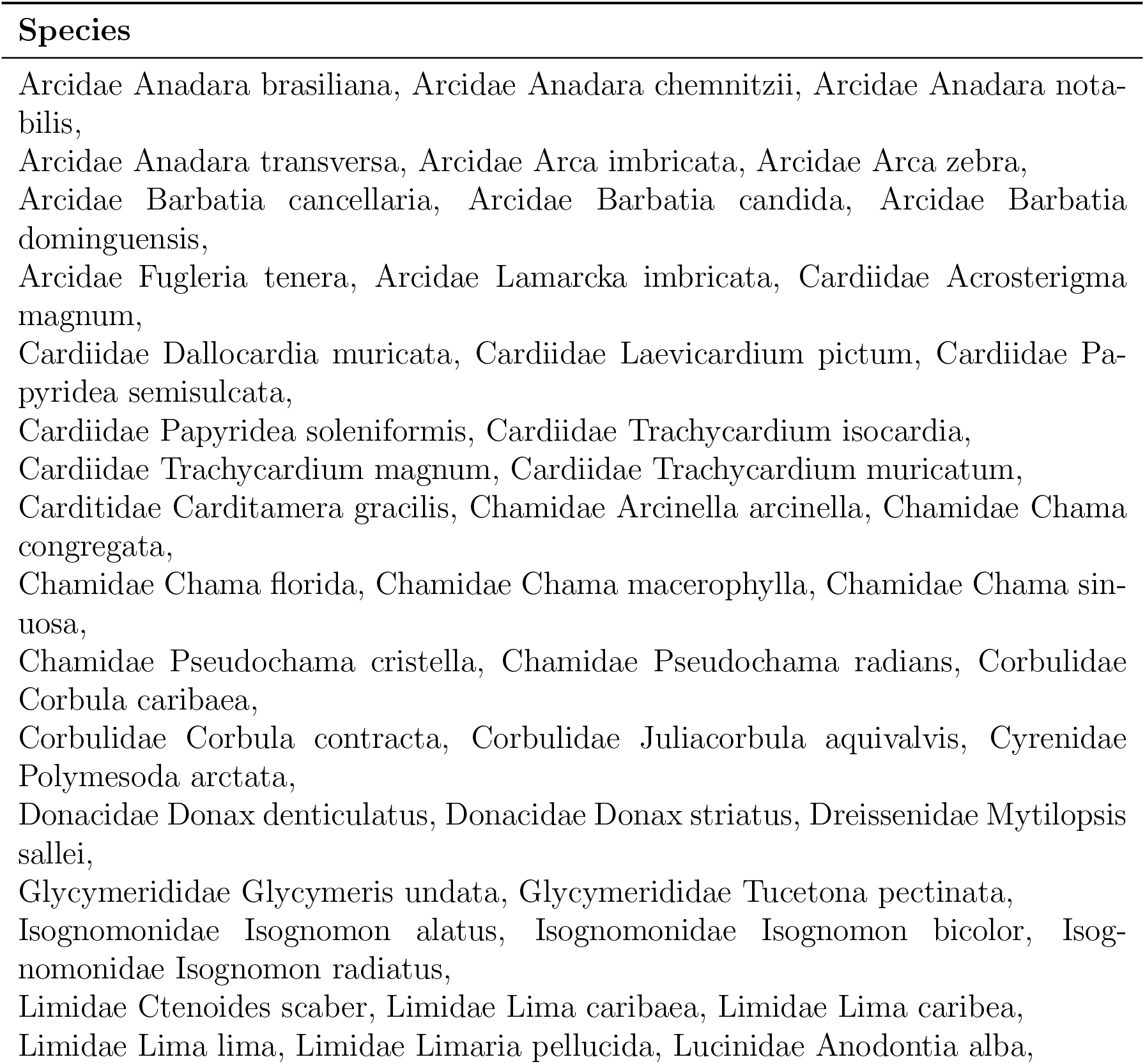
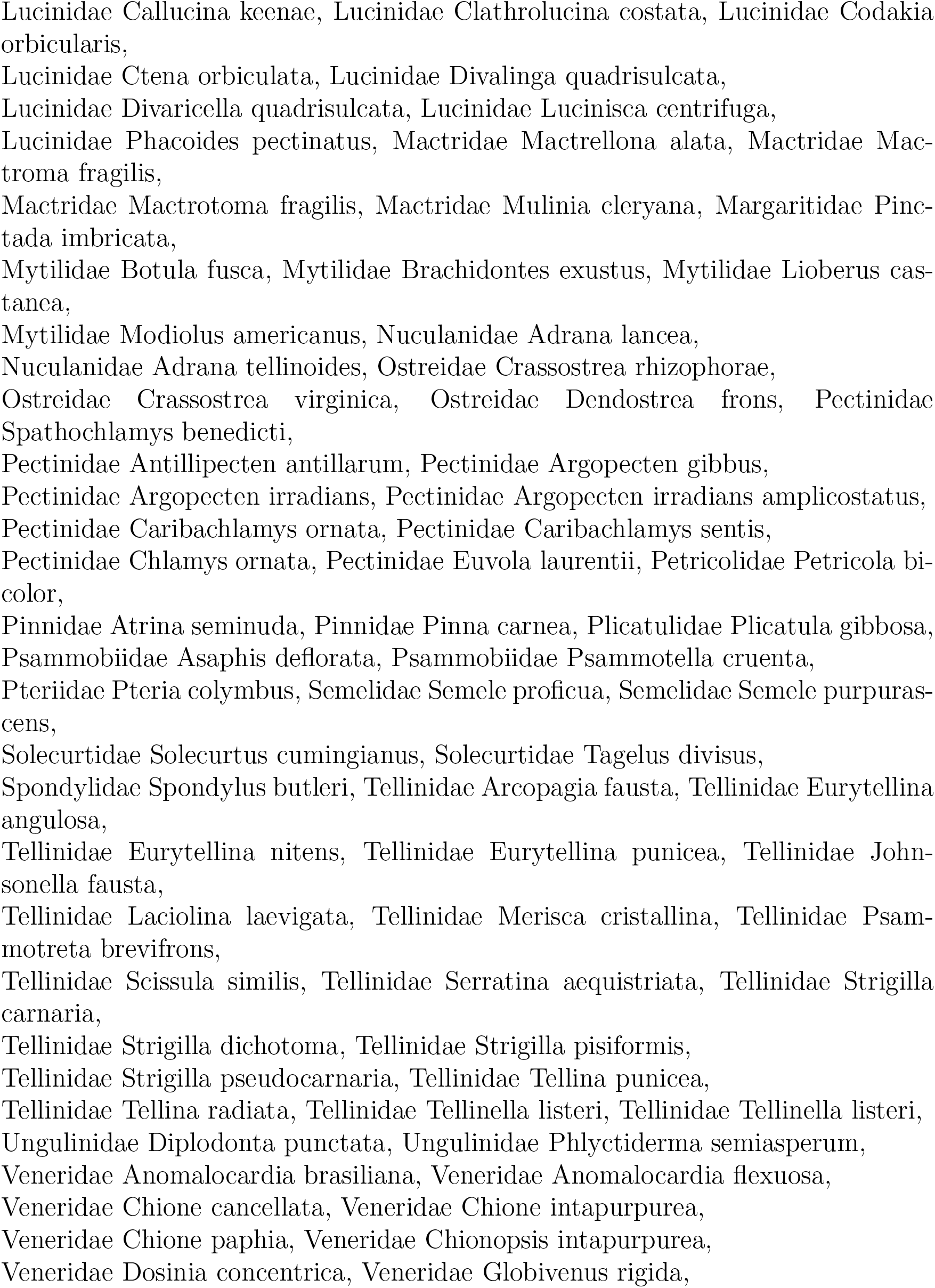
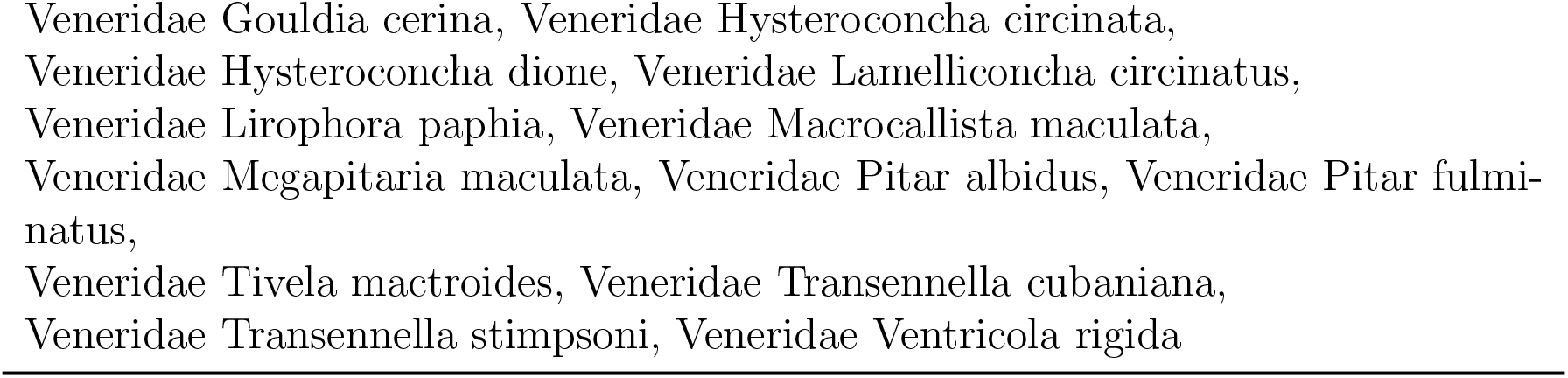
Seashell Species - Caribbean Bivalves.

**Table 9:**
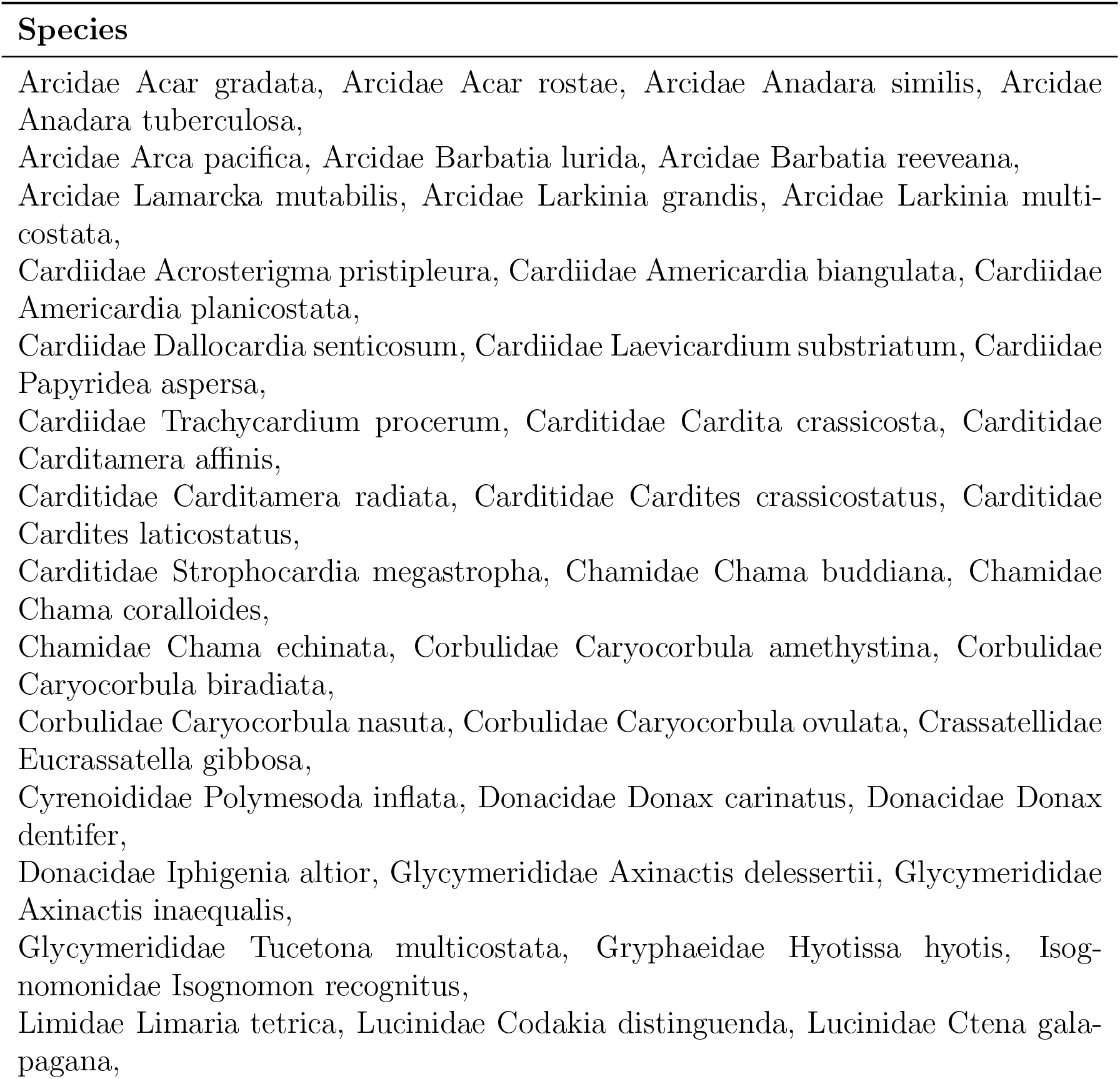
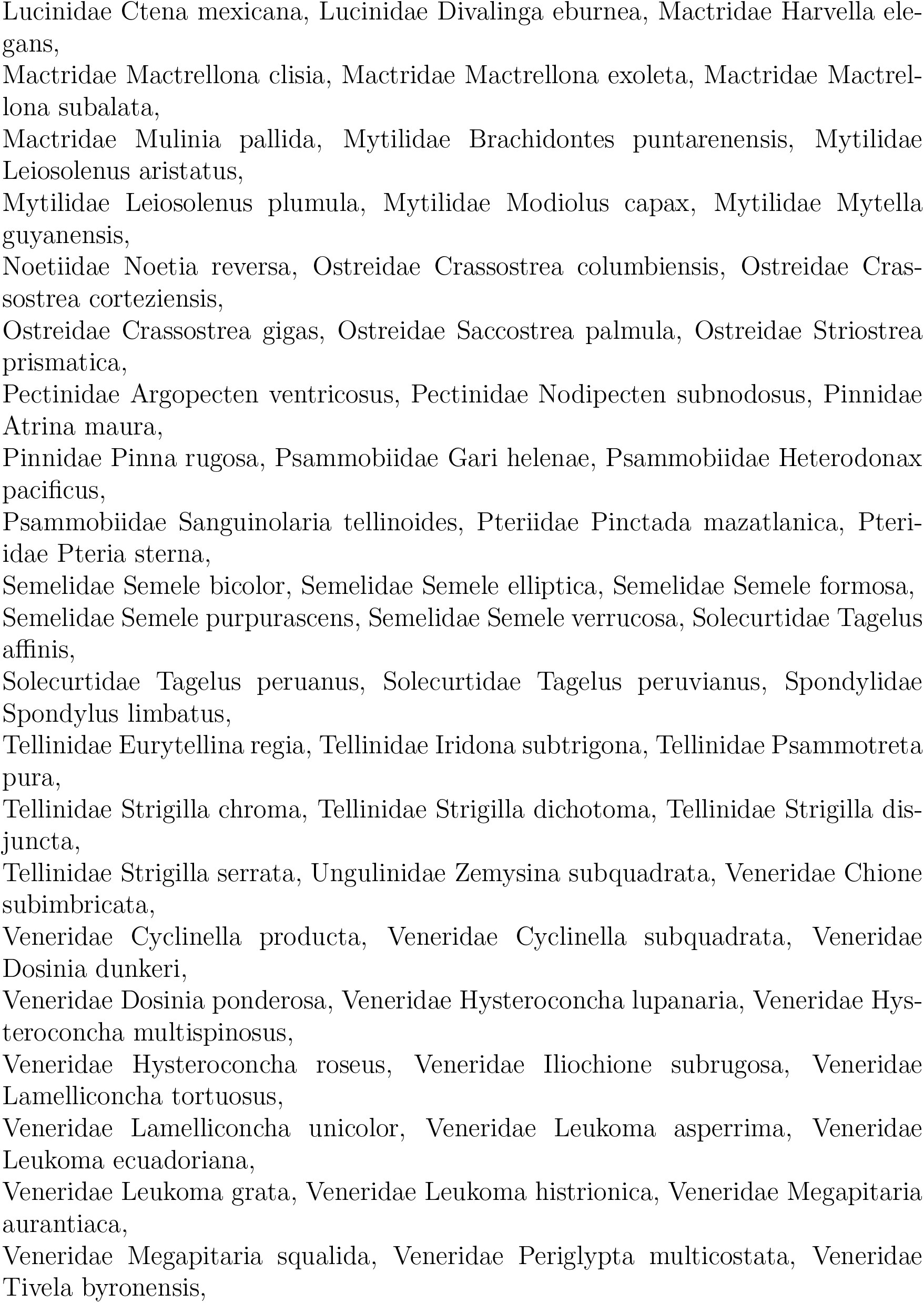
Seashell Species - Pacific Bivalves.

